# Multiplexed Ultrasound Imaging of Gene Expression

**DOI:** 10.1101/2024.10.30.621148

**Authors:** Nivin N. Nyström, Zhiyang Jin, Marisa E. Bennett, Ruby Zhang, Margaret B. Swift, Mikhail G. Shapiro

## Abstract

Acoustic reporter genes (ARGs) have enabled the imaging of gene expression with ultrasound, which provides high-resolution access to deep, optically opaque living tissues. However, unlike their fluorescent counterparts, ARGs have so far been limited to a single “color,” preventing multiplexed imaging of cellular states or populations. Here, we use rational protein design and directed evolution to develop two novel ARGs that can be distinguished from each other based on their acoustic pressure response profiles, enabling “two-color” ultrasound imaging of gene expression. We demonstrate the utility of multiplexed ARGs for delineating bacterial cell species and cell states *in vitro*, and then apply them towards imaging distinct subpopulations of probiotics in the mouse gastrointestinal tract and in tumor-colonizing bacterial agents *in vivo*. Just as the first wavelength-shifted derivatives of fluorescent proteins opened a vivid world for optical microscopy, our next-generation acoustic proteins set the stage for a richer symphony of ultrasound signals from living subjects.

## INTRODUCTION

Fluorescent reporter genes have revolutionized the study of living systems. Since the discovery of GFP, an assortment of spectrally orthogonal fluorescent proteins have been developed and are now routinely employed for multiplexed imaging of distinct molecular events within a single field of view.^1-3^ While fluorescence offers unparalleled spatial resolution and sensitivity, it is limited to *ex vivo* samples or superficial tissues *in vivo* (≤1 mm) due to light scattering and attenuation.^4^

For deep tissue imaging in living subjects, ultrasound leverages the favorable interactions of acoustic waves with tissue to permit routine imaging at depths of several centimeters and resolutions on the order of 100 µm. In addition, ultrasound imaging offers portability, low cost, and high temporal resolution.^5^ Recently, acoustic reporter genes (ARGs) that encode air-filled protein nanostructures called gas vesicles (GVs) have been established as “acoustic proteins” for ultrasound imaging of cellular function. However, these acoustic analogs of GFP have so far been limited to a single “color” readout.^6-8^

We wondered whether it might be possible to generate additional “colors” of ARGs to enable multiplexed imaging, analogous to the different varieties of fluorescent proteins. This would allow us to measure multiple variables simultaneously within a single experiment, increasing throughput and data dimensionality, which currently stand as major bottlenecks in contemporary *in vivo* imaging methods.^9^

To enable such multiplexing, we sought to engineer ARGs with unique pressure responses. We focused on engineering the secondary structural protein GvpC that scaffolds the GV shell and modifies its stiffness. GVs with a softer shell undergo reversible mechanical buckling under acoustic pressure waves, resulting in nonlinear ultrasound contrast.^10-12^ We hypothesized that by engineering GvpC we could tune the buckling pressure threshold of the GVs, thereby producing variants with different pressure thresholds, allowing us to tell them apart by imaging with a series of suitable pressures. To test this hypothesis, we designed a high-throughput *in cellulo* acoustic screen and used it to engineer and evolve two next-generation ARGs, which we call bARG_560_ and bARG_710_, to multiplex gene expression on ultrasound images.

To demonstrate the utility of these ARGs, we introduced them into probiotic microorganisms, allowing us to delineate two species – *Salmonella typhimurium SL1344* (Stm) and *E. coli Nissle 1917* (EcN) – within a single field-of-view. Critically, unlike previous attempts at acoustic multiplexing,^6,10^ distinguishing bARG_560_ from bARG_710_ did not require irreversibly collapsing the GVs and thereby destroying their ultrasound contrast. We also incorporated bARG_560_ and bARG_710_ into a genetic state-switch circuit to visualize two cellular states with ultrasound. Finally, we employed these new ARGs for multiplexed, deep tissue, high spatial resolution imaging in live mice, demonstrating the capacity to distinguish between microbial species in the gastrointestinal tract and track co-colonization of tumors by distinct populations of tumor-homing bacteria.

## RESULTS

### Removal of *gvpC* and construct optimization results in ARGs with a low pressure threshold and strong nonlinear contrast

To develop multiplexed ARGs, we started with the most recently developed bacterial ARG system derived from *Serratia sp. 39006* (bARG_Ser_), which has been used to enable sensitive nonlinear imaging of genetically modified bacteria *in vitro* and *in vivo*.^8^ Alongside other GV-forming genes, bARG_Ser_ encodes GvpC, the alpha-helical protein that binds to the GV shell to increase its stiffness. To generate ARGs with more permissive shell buckling, and therefore nonlinear contrast at lower acoustic pressures, we first deleted *gvpC* from bARG_Ser_ and replaced it with *gfp* (**Fig. 1a**), and used a high-throughput optical assay, based on the optical scattering of assembled GVs in multiwell bacterial patches, to optimize the genetic construct and its expression conditions (**Fig. 1b-c**). Screening ∼10^4^ unique conditions (**Fig. 1c**) yielded a Δ*gvpC* construct with highly efficient expression and minimal burden. In this construct, the GV operon is transcribed by an IPTG-dependent T7 RNA polymerase expressed under L-arabinose control (**Supplementary Table T1**).

**Figure 1.**
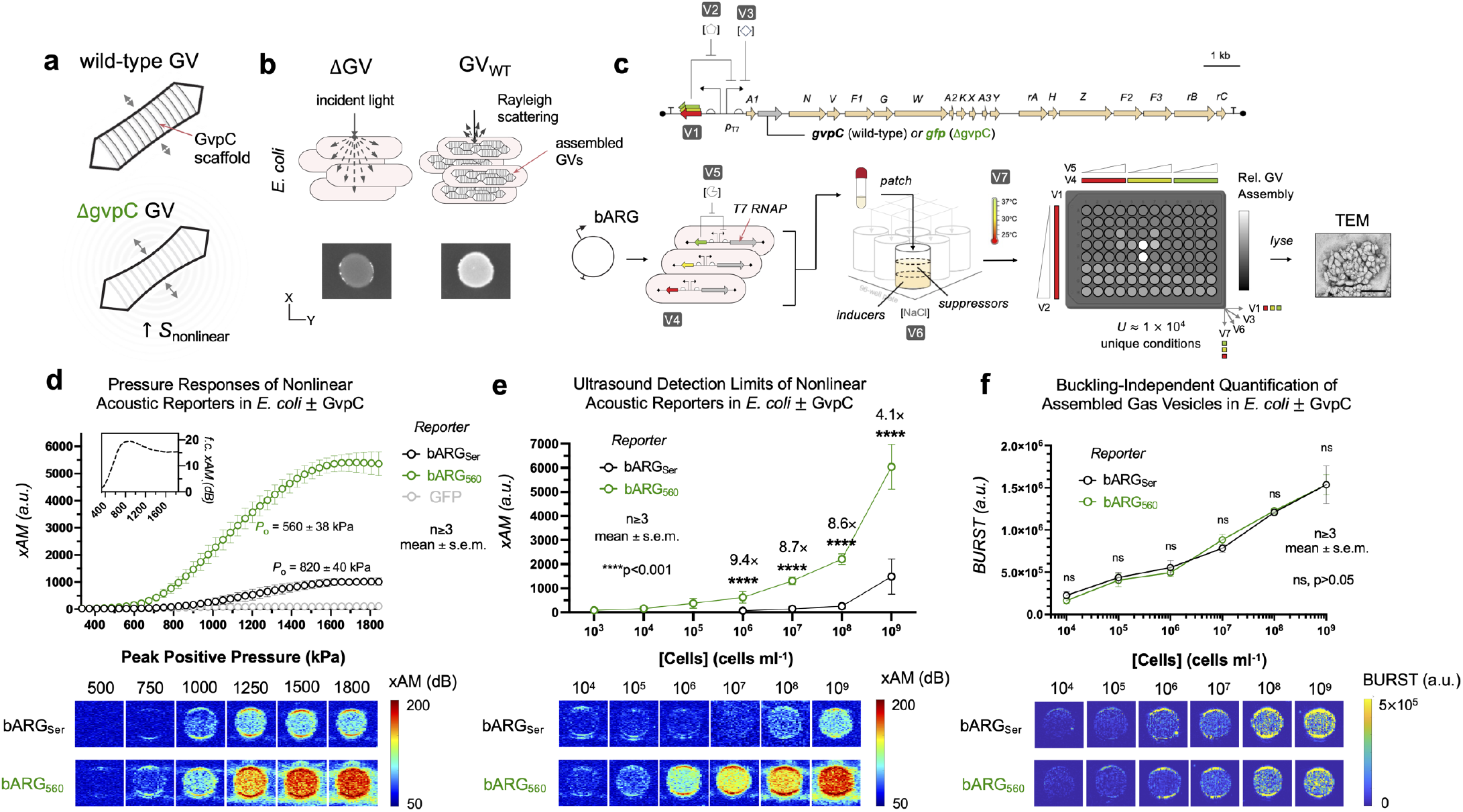
*In cellulo* assembly screening for an acoustic reporter gene with increased nonlinearity. **(a)** The wild-type gas vesicle (GV) nanostructure with its native GvpC protein scaffold (*top*). GVs lacking GvpC exhibit an increased amplitude of reversible buckling resulting in stronger nonlinear ultrasound signal (*bottom*). **(b)** GVs assembled within *E. coli* scatter light, resulting in patches with increased opacity. **(c)** A high-throughput assembly screen to identify circuit conditions for *in cellulo* assembly of GVs with a *gvpC* deletion. *Top*: The annotated *Serratia sp. 39006* GV operon, with *gvpC* (bARG_Ser_), or with *gvpC* replaced by *gfp* (bARG_560_). Scale bar, 1 kb. *Bottom, from left to right*: plasmids were transformed into *E. coli* and grown in liquid culture, before being patched on dual-layer 96-well agar plates. Plates were assessed for GV assembly via patch opacity. TEM, transmission electron microscopy. Scale bar, 200 nm. **(d)** Nonlinear acoustic signals measured at variable pressures from *E. coli* encoding bARG_Ser_ or bARG_560_ (mean ± s.e.m., n≥4). xAM, X-wave amplitude modulation signal. *Inset*: Nonlinear signal fold-change (f.c.) across pressures between bARG_Ser_ and bARG_560_ in decibels (dB). **(e)** Nonlinear acoustic signals of *E. coli* encoding bARG_Ser_ or bARG_560_ at variable cell concentrations (mean ± s.e.m., n≥3). f) Buckling-independent quantification of GVs in *E. coli* encoding bARG_Ser_ or bARG_560_ at variable cell concentrations (mean ± s.e.m., n≥3).

To measure the resulting acoustic contrast, we loaded cells into agarose phantoms and acquired nonlinear ultrasound images at transmit pressures ranging from 0.3 to 1.8 MPa using the xAM amplitude modulation pulse sequence^13^ (xAM; n≥3). With our new construct, we observed a decreased nonlinear threshold (*P*_o_ = 560 ± 38 kPa) and an increased nonlinear signal yield (*S*_max_ = 5900 ± 660 a.u.) relative to the wild-type operon (*P*_o_ = 820 ± 40 kPa, *S*_max_ = 1000 ± 130 a.u.; **Fig. 1d**). We named this new construct bARG_560_ based on its nonlinear pressure threshold. With respect to imaging sensitivity, bARG_560_ exhibited 4-to 9-fold higher nonlinear signals relative to bARG_Ser_ across a range of cell concentrations (**Fig. 1e**), while overall GV production, as measured using the buckling-independent BURST pulse sequence,^14^ was not significantly different between the two populations (n≥3, p>0.05, **Fig. 1f**).

### Directed evolution of GvpC results in ARGs with shifted pressure response profiles

To engineer a second ARG with strong nonlinear scattering and a pressure threshold distinctly higher than that of bARG_560_, we performed a directed evolution screen of GvpC (**Fig. 2a**). Our target pressure profile would result a nonlinear signal threshold upshifted by 100 kPa or more relative to bARG_560_, while producing strong contrast below the collapse pressure of bARG_560_ (*P*_c_ = 1220 ± 34 kPa, **Supplementary Fig. S1**) to enable continuous multiplexed imaging without destroying GVs. To achieve these specifications, we ran a directed evolution campaign (**Fig. 2a**). We used error-prone PCR to generate a pooled mutagenesis library of *gvpC*, induced *E. coli* colonies containing the resulting variants on solid agar, and loaded samples into 96-well agarose phantoms for acoustic measurements on a custom-built motorized ultrasound system (**Fig. 2a**). To organize our library dataset, we first set a linear scattering (B-mode) intensity threshold (*S*_B_ ≥ 175 a.u.) to filter out samples with insufficient GV expression, and plotted all remaining mutants according to their nonlinear signal threshold (*P*_o_) and maximum nonlinear signal yield (*S*_max_; **Fig. 2b**). Altogether, we obtained data from 880 variants.

**Figure 2.**
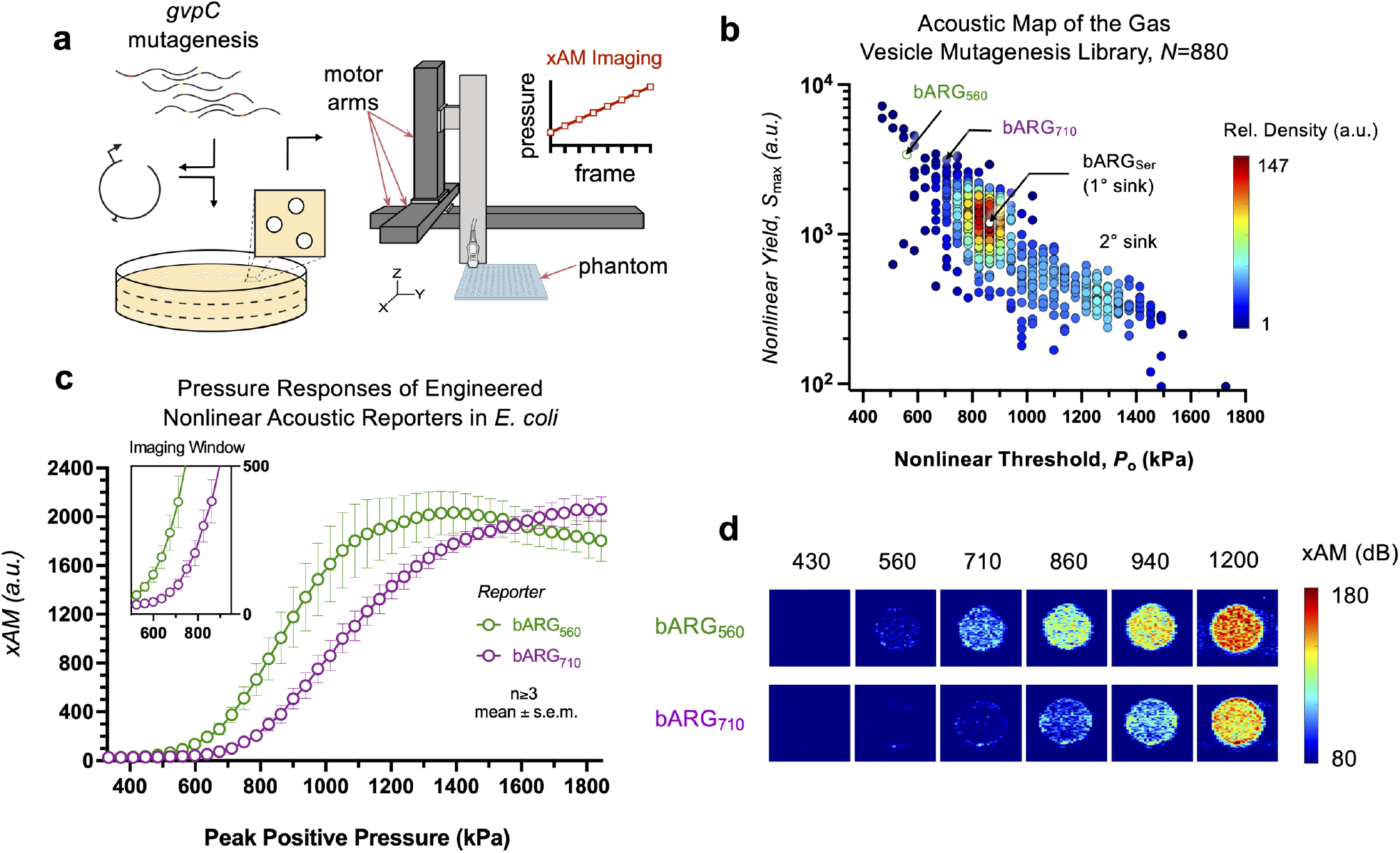
Directed evolution of acoustic reporter genes for pressure-domain multiplexing. **(a)** An error-prone PCR library of *gvpC* was cloned into a backbone plasmid containing the bARG_560_ operon, transformed into *E. coli* and then plated. Colonies were arrayed, grown and induced for expression, after which, they were loaded into agarose phantoms for nonlinear acoustic measurements on a custom-built motorized ultrasound system. **(b)** Acoustic map of the *gvpC* mutagenesis library. *P*_o_, nonlinear pressure threshold. *S*_max_, maximum nonlinear yield. Each point represents a mutant (*N*=880). Color scale represents the relative density of mutants on the plot. Controls include bARG_Ser_ (*white circle*) and bARG_560_ (*green circle*). **(c)** Nonlinear acoustic signals measured at increasing pressures from *E. coli* encoding bARG_560_ and bARG_710_ (mean ± s.e.m., n≥3). xAM, *Inset*: Nonlinear imaging window, capped at the collapse pressure of bARG_560_. **(d)** Nonlinear images of *E. coli* encoding bARG_560_ or bARG_710_ acquired across increasing pressures. dB, decibels.

We observed a strong negative correlation between the nonlinear threshold and maximum nonlinear yield of the variants in our library, which was not unexpected: *gvpC* sequences that confer greater rigidity to the GV shell require larger force to initiate buckling are also likely to limit the amplitude of GV deformation (and resulting sound scattering) after buckling. bARG_560,_ included as a control, sits near the upper left corner of the acoustic map (*P*_o_ = 560 kPa, *S*_max_ = 3400 a.u.). We observed two acoustic “sinks” – regions of the plot that contained high concentrations of sequences. The deeper sink was centered on the parent *gvpC* (bARG_Ser_, *P*_o_ = 860 kPa, *S*_max_ = 1190 a.u.) from which all these mutants were derived. The second and shallower sink contained sequences with relatively high nonlinear thresholds and low nonlinear signal within the tested pressure range.

To gain a deeper understanding of our sequence landscape and how it relates to our acoustic map, we sequenced a subset of mutants from different acoustic regions (1° sink, 2° sink, the bridge between them, and outliers that resided outside of these regions), and performed pairwise Needleman-Wunsch protein sequence alignments of GvpC against the wild-type parent (**Supplementary Fig. S2**). Not surprisingly, we found that sequences within the 1° acoustic sink exhibited the lowest sequence distance scores (*D*_1_ = 0.067 ± 0.006, *N*=57), signifying that single mutations were generally conservative with respect to acoustic properties. We saw a significant increase in sequence distance in variants in the 2° sink (*D*_2_ = 0.48 ± 0.04, *N*=28) and the bridge between the sinks (*D*_b_ = 0.39 ± 0.04, *N*=79). Most notably, we found that sequences classified as acoustic outliers exhibited the highest sequence distance scores on average (*D* = 0.58 ± 0.04, *N*=85).

After evaluating several outlier variants that met our design specifications for pressure-domain multiplexing, we selected a mutant containing an L154P mutation that conferred a nonlinear signal threshold of 710 ± 32 kPa and a nonlinear yield comparable to that of bARG_560_ within the “non-destructive imaging window” below 1200 kPa (**Fig. 2c**). An AlphaFold_15_ structural prediction of GvpC_L154P_ compared to the parent protein suggests that a proline-imposed kink in the alpha helix may result in reduced contact with the GV shell (**Supplementary Note 1, Supplementary Fig. S3**). We call this evolved construct bARG_710_, according to its nonlinear threshold in kPa.

### Multiplexed ultrasound can distinguish two cell types or cell states

After obtaining two ARGs with acoustic properties compatible with multiplexing, we tested their ability to render two cell types or two cell states visible to ultrasound in the same sample. The ability to delineate distinct cell populations in deep tissues of living subjects at high spatial resolution would be useful across many applications, including cell-based diagnostics and therapies.^16,17^ As a first step in this direction, we engineered two widely studied probiotic microorganisms – attenuated *Salmonella typhimurium SL1344* (Stm)^18^ and *E. coli Nissle 1917* (EcN)^19^ – to express bARG_560_ and bARG_710_, respectively These probiotic strains are widely used in the development of cell-based diagnostics and therapies due to their ability to populate the gastrointestinal tract, colonize the hypoxic core of tumors, sense local biomarkers, and produce therapeutic payloads, and have already entered human clinical trials.^20-22^

To implement our new ARGs in these two species, we extensively optimized the corresponding genetic constructs by testing several *gvp* gene deletions and promoter sequences (**Supplementary Note 2; Supplementary Fig. S4, Supplementary Fig. S5**) on an axe/txe stabilized plasmid.^23^ For bARG_710_, we placed *gvpC*_L154P_ onto its own operon under regulation of a constitutive promoter derived from *pTac*,^24^ and co-expressing a red fluorescence reporter gene, *dsRed2*.

To enable the acquisition of multiplexed ultrasound images, we generated a multiplexing matrix (***M***) by measuring nonlinear signals from pure populations of bacteria encoding either bARG_560_ or bARG_710_ across the pressure domain. We then used ***M*** for pixelwise linear unmixing of images containing our samples, solving for the coefficient matrix (***C***) of our images via simple linear algebra (**Fig. 3a; Supplementary Table T3**). ***C*** contains the estimated contributions of our two “acoustic channels” to the signal in each pixel, allowing us to create spatial maps of each reporter gene.

**Figure 3.**
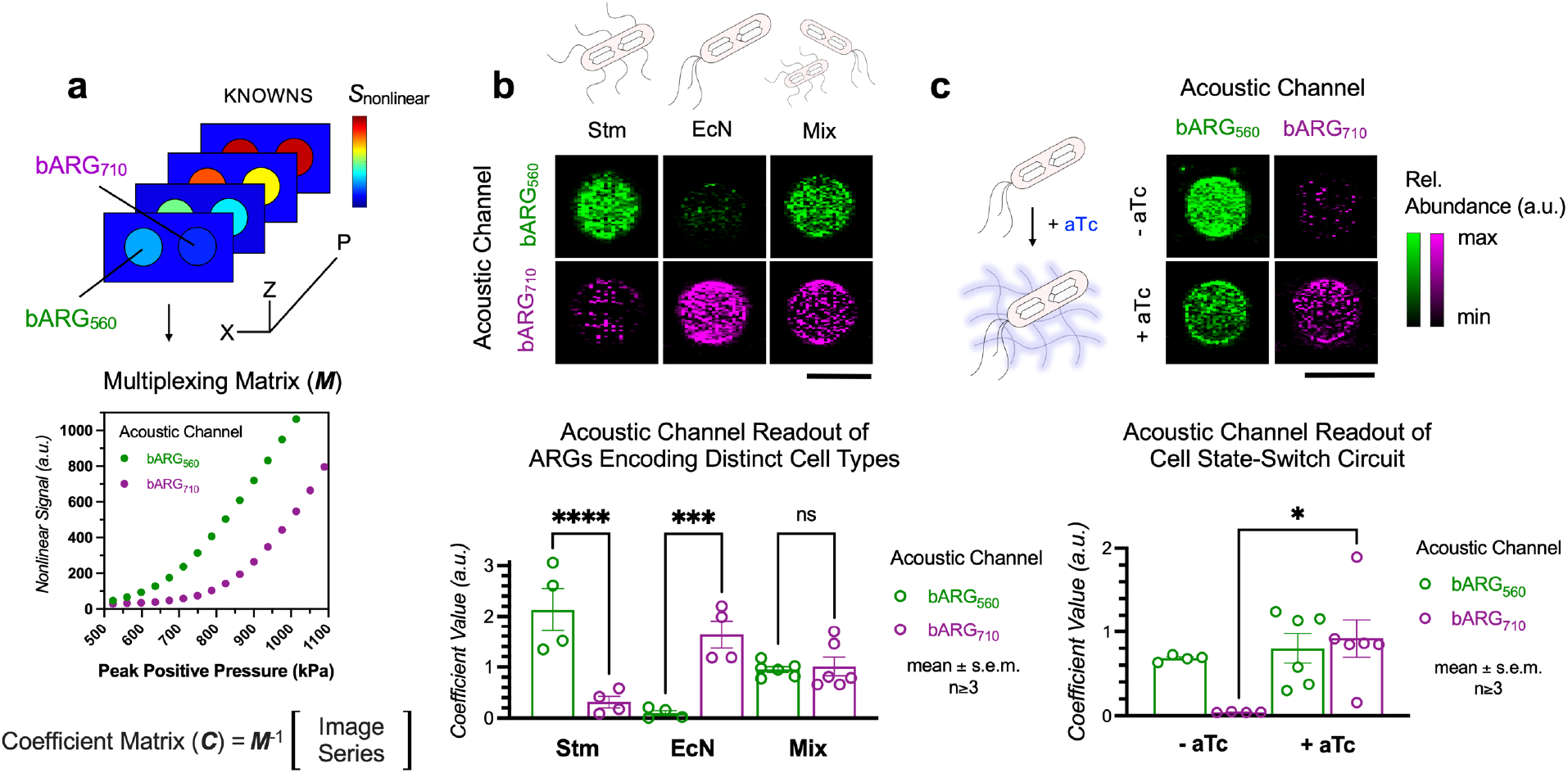
Multiplexed imaging cell types and cell states on ultrasound with acoustic reporter genes. **(a)** A multiplexing matrix ***M*** was produced by measuring nonlinear signals from pure populations of cells encoding bARG_560_ or bARG_710_ across the pressure domain. ***M*** was then used for pixelwise linear unmixing of input images containing pure or mixed reporter samples, solving for the coefficient matrix ***C*. (b)** Representative coefficient matrix images, or “acoustic channel” images, of pure or mixed populations of attenuated *Salmonella typhimurium SL1344* (Stm) encoding bARG_560_ and *E. coli Nissle 1917* (EcN) encoding bARG_710_. Scale bar, 2 mm. Mean coefficient values (mean ± s.e.m., n≥3, a.u.) of acoustic channel images from pure or mixed populations of Stm encoding bARG_560_ and EcN encoding bARG_710_. **(c)** Representative acoustic channel images of cells encoding the tetracycline-responsive bARG_560_/bARG_710_ state-switch circuit, induced for expression in the absence (-) or presence (+) of anyhydrotetracycline (aTc). Scale bar, 2 mm. **(d)** Mean coefficient values (mean ± s.e.m., n≥3, a.u.) of acoustic channel images from cells induced for expression in the absence or presence of aTc.

As expected, in pure samples of each type, we observed signals almost entirely on the bARG_560_ channel in images of Stm encoding bARG_560_ (*S*_Stm/560_ = 2.1 ± 0.4 a.u., *S*_Stm/710_ = 0.31 ± 0.1 a.u., p<0.0001) and the reverse relationship for images of EcN encoding bARG_710_ (*S*_EcN/560_ = 0.099 ± 0.05 a.u., *S*_EcN/710_ = 1.64 ± 0.3 a.u., p=0.0007). In images of a mixture of Stm and EcN, we observed similar signals from the two channels (*S*_mix/560_ = 0.95 ± 0.06 a.u., *S*_mix/710_ = 1.01 ± 0.2 a.u., **Fig. 3b**).

In addition to multiplexing separate cell populations, we wanted to enable the imaging of distinct cell states in a single population, for example ones triggered by an environmental cue. As a proof-of-concept, we built a construct which by default expresses the genes encoding bARG_560_, but switches to bARG_710_ in response to tetracycline (aTc) by expressing *gvpC*_L154P_ (**Supplementary Fig. S6**). When we expressed this circuit in EcN cells and imaged them with ultrasound, we found that addition of aTc led to a large increase in signal on the bARG_710_ channel (*S*_-aTc/710_ = 0.042 ± 0.002 a.u., *S*_+aTc/710_ = 0.92 ± 0.2 a.u., p=0.02; **Fig. 3b**).

### Multiplexed ARGs enable the imaging of distinct cell populations in the gastrointestinal tract

To provide a technical demonstration of multiplexed ultrasound imaging in the context of living tissue, we imaged our two ARG-expressing bacterial species (Stm-bARG_560_ and EcN-bARG_710_) in the gastrointestinal tract of live mice. To have precise control over their *in vivo* distribution, we transplanted these cells into the mouse colon using hydrogel injection. This model has served as a well-controlled technical proving ground in previous studies of reporter genes and biosensors for ultrasound.^6,25^ Here, we transplanted the cells either as pure populations or as mixed communities (**Fig. 4a**). We used B-mode imaging to locate a cross-section of the gastrointestinal lumen based on soft tissue contrast before acquiring xAM images across the pressure domain. We then performed linear unmixing on the pressure-domain image datasets to generate two coefficient matrices for each mouse, representing each acoustic channel (**Fig. 4b**).

**Figure 4.**
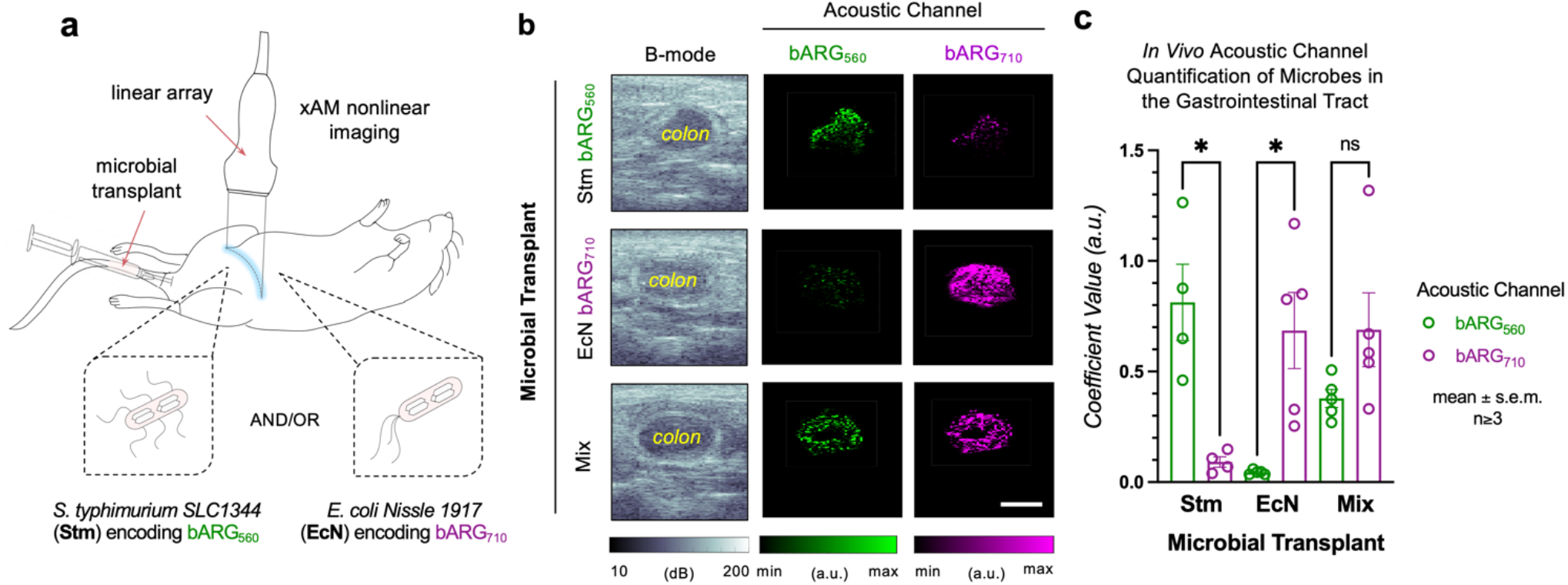
Imaging distinct microbial cell populations in guts of living mice on ultrasound using nonlinear acoustic reporter genes. **(a)** Microbial transplant model. Attenuated *S. typhimurium SL1344* (Stm) encoding bARG_560_ and *E. coli Nissle 1917* (EcN) encoding bARG_710_ were transplanted into the gastrointestinal system of mice. B-mode imaging and xAM nonlinear imaging was performed across the pressure domain, followed by a high pressure collapse pulse. **(b)** Representative B-mode images, acoustic channel images, and merged overlay. Decibels, dB. Arbitrary units, a.u. Scale bar, 2 mm. **(c)** Mean coefficient values (mean ± s.e.m., n≥3, a.u.) of acoustic channel images of the gastrointestinal lumen of mice that received microbial transplants of Stm encoding bARG_560_ and/or EcN encoding bARG_710_.

We saw that mice transplanted with Stm-bARG_560_ generated significantly greater mean coefficient values in the bARG_560_ channel (*S*_Stm/560_ = 0.81 ± 0.2 a.u.) than the bARG_710_ channel (*S*_Stm/710_ = 0.091 ± 0.02 a.u., p=0.02, n=4). Similarly, we saw that mice transplanted with EcN-bARG_710_ generated significantly greater mean coefficient values in the bARG_710_ channel (*S*_EcN/710_ = 0.69 ± 0.2 a.u.) than the bARG_560_ channel (*S*_EcN/560_ = 0.044 ± 0.004 a.u., p=0.02, n=4). Finally, we observed significantly increased mean coefficient values in both the bARG_560_ (*S*_mix/560_ = 0.38 ± 0.04 a.u.) and bARG_710_ (*S*_mix/710_ = 0.69 ± 0.2 a.u.) channels from GI images of mice that harbored both microbial populations, relative to signals from negative acoustic channels in control mice (**Fig. 4c**). These experiments validated that despite the greater aberration, attenuation and background contrast of living tissues, pressure-based acoustic multiplexing successfully distinguishes two “colors” of ARGs.

### Multiplexed ARGs enable the imaging of tumor homing by two distinct tumor-colonizing bacteria

Having provided a technical demonstration for *in vivo* multiplexed imaging in the GI tract, we sought to demonstrate the utility of acoustic multiplexing in a translationally relevant application by imaging two tumor-colonizing bacterial species in the same therapy recipient. Recent research has established that tumors are naturally colonized by bacterial species,^26,27^ and that certain strains of bacteria can be used as tumor-colonizing therapeutic agents due to their ability to thrive in the immunosuppressed, hypoxic, necrotic cores of tumors and be engineered to produce oncolytic or immunostimulatory biologics to induce remission.^20,22^ Within this paradigm, ultrasound can offer the ability to track the successful engraftment of a probiotic cell therapy at its target, while multiplexing could provide a way to track successively administered doses, co-administered variants, or conditional cellular states.

To provide a proof of concept for multiplexed cell therapy imaging in tumors, we implanted murine colon cancer cells into the right flanks of mice and allowed the tumors to grow for 14 to 21 days before intravenously injecting pure or mixed populations of Stm-bARG_560_ and EcN-bARG_710_ (**Fig. 5a**). We induced GV expression for three days, starting 24 h after injecting the bacteria, and performed ultrasound imaging on the fourth day. We observed dominant signal from the bARG_560_ channel compared to the bARG_710_ channel in tumors of mice injected with Stm (*S*_Stm/560_ = 0.71 ± 0.2 a.u., *S*_Stm/710_ = 0.026 ± 0.002 a.u., p=0.006, n=4) and significantly higher signal from the bARG_710_ channel relative to the bARG_560_ channel in tumors of mice injected with EcN (*S*_EcN/560_ = 0.063 ± 0.02 a.u., *S*_EcN/710_ = 0.58 ± 0.2 a.u., p=0.01, n=5).

**Figure 5.**
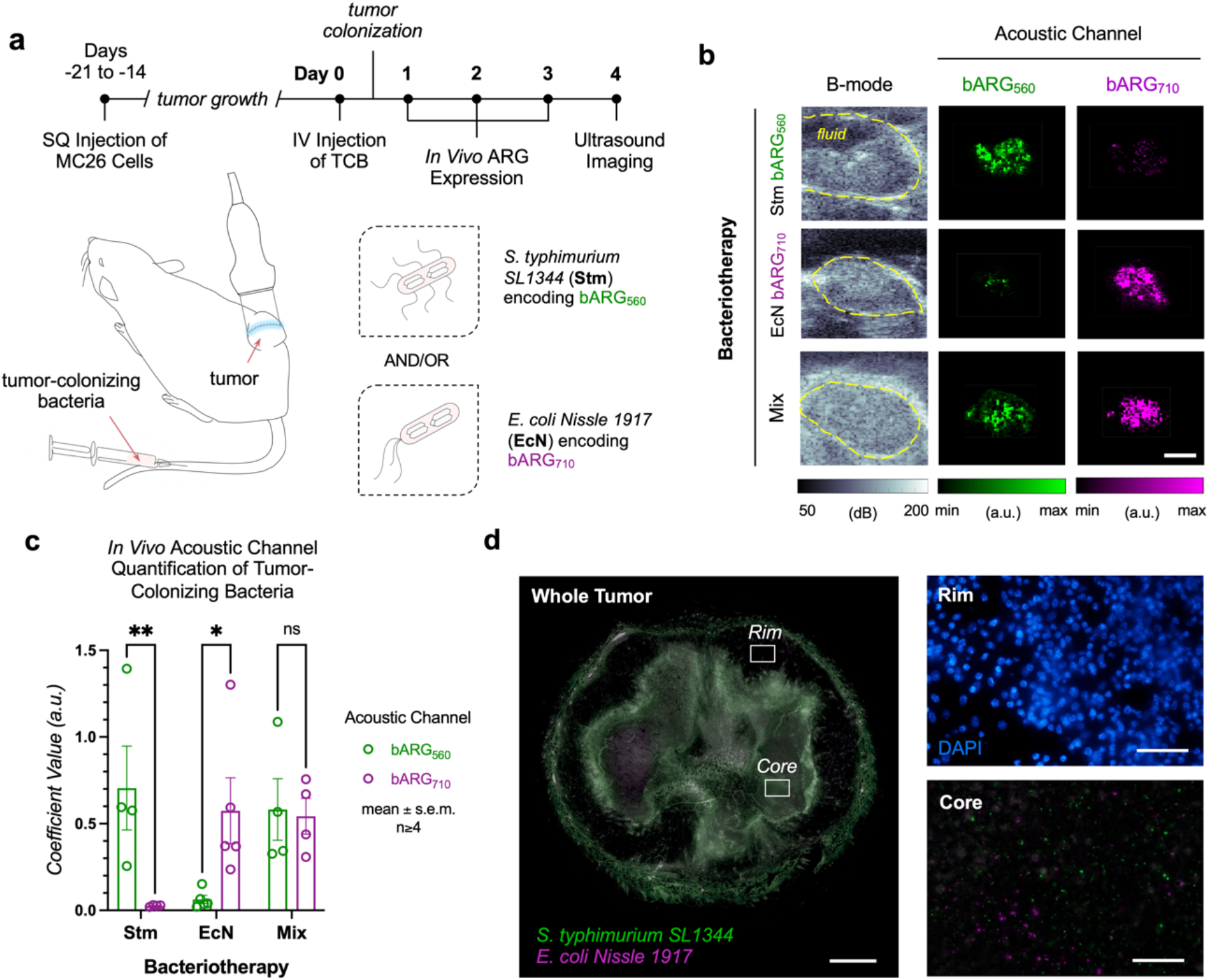
Multiplexed ultrasound imaging of tumor colonization by single- or dual-species probiotic agents. **(a)** Mice received subcutaneous injections of MC26 tumor cells into their right hind flanks. Once tumors grew, a dose of tumor-colonizing bacteria, consisting of either attenuated *S. typhimurium SL1344* (Stm) encoding bARG_560_ and/or *E. coli Nissle 1917* (EcN) encoding bARG_710_, were intravenously injected and given 24 h to colonize the tumor. The following day, and every 24 h for the following two days, mice were intraperitoneally injected with L-arabinose to induce GV expression *in situ*. B-mode images and xAM images across the pressure domain were then acquired to image the distribution of each species. **(b)** Representative B-mode images and nonlinear bARG_560_ (*green*) and bARG_710_ (*magenta*) acoustic channel images of mice receiving different species or mixed populations of tumor-colonizing bacteria. Decibels, dB. Arbitrary units, a.u. Scale bar, 2 mm. **(c)** Mean coefficient matrix values (a.u.) of bARG_560_ (*green*) and bARG_710_ (*magenta*) acoustic channels from tumors receiving tumor-colonizing bacteria communities consisting of Stm encoding bARG_560_ and/or EcN encoding bARG_710_. **(d)** Whole-tumor fluorescence image overlay of GFP *(green)*, and dsRed2 (*magenta*) channels from a representative mouse that received a 1:1 species mixture (v/v) of attenuated *S. typhimurium SL1344* encoding bARG_560_ (*green*) and *E. coli Nissle 1917* encoding bARG_710_ (*magenta*). White regions represent overlays of both species. Scale bar, 1 mm. High-magnification fluorescence images of the tumor core and the tumor rim. Scale bar, 100 µm.

When we administered a 1:1 mixture of Stm and EcN, we observed comparable levels of signals from each acoustic channel (*S*_mix/560_ = 0.58 ± 0.2 a.u., *S*_mix/710_ = 0.54 ± 0.1 a.u., p=0.85, n=4; **Fig. 5b-c**), confirming the ability to distinguish pure from mixed colonization scenarios. To validate that Stm and EcN both colonized the tumor in mice that received mixed population doses, and that interspecies competition did not result in single-strain dominance, we performed tumor histology. In agreement with our ultrasound images, we saw co-colonizatoin of the tumor core by both bacterial species, and validated *in vivo* induction of both bARG_560_ and bARG_710_ operons through visualization of their fluorescent markers within the same tumor (**Fig. 5d**).

## DISCUSSION

This work establishes multiplexed ultrasound imaging of gene expression using two next-generation ARGs. We anticipate that the ability to track multiple cell types or states simultaneously within individual living subjects will enable more comprehensive study of complex biological systems. While other imaging modalities have already benefited from multiplexing,^28-30^ ultrasound can now do so for the first time.

In the process of engineering bARG_560_ and bARG_710_, we established bespoke methods for ARG engineering that may be extended to related applications. For example, the high-throughput genetic optimization we used to enable bARG_560_ expression could help domesticate GV-encoding gene clusters from other species or adapt ARGs to additional microbial species such as native members of the gut microbiome.

We anticipate several improvements and extensions to this technology. One limitation of acoustic pressure-based multiplexing is that it requires the ability to apply controlled pressure levels within tissues. While we had no difficulty doing so in mouse tumors and colons, it may be more challenging in more attenuating scenarios. To address such cases, it may be necessary to develop pulse sequences that estimate attenuation aberration and correct for it by shaping transmitted pulses.

An important extension of the technology would be to implement acoustic multiplexing in mammalian ARGs, which are being used to image a variety of cell types *in vivo*, including tumors and immune cells. The presence of *gvpC* in mammalian ARGs suggests that an approach analogous to the one taken here could uncover pressure-shifted variants, although they will likely need to be screened in the mammalian context.

Another valuable extension would add more ARG “colors” to enable higher-order multiplexing. Just as fluorescent proteins have been genomically mined, evolved and engineered for decades to generate an array of reporter genes for multiplexed imaging, which have since been used in innumerable optical imaging applications, the methods and constructs developed here suggest that a similar trajectory may be available for ultrasound.

## MATERIALS AND METHODS

### Molecular Biology

For all genetic constructs, primers were manually designed and ordered from Integrated DNA Technologies (IDTs), and all enzymes were ordered from New England Biolabs (NEB), unless otherwise stated. The constitutive promoter used in our circuits was a gift from Howard Shuman (Addgene plasmid #84821). Polymerase chain reactions (PCRs) were amplified using Taq (M0270, NEB) for fragments <3 kb and Q5 (M0492, NEB) for fragments >3 kb, and Gibson assembly was carried out using the HiFi DNA Assembly Master Mix (E2621, NEB). All assembly reactions were transformed into Stable *E. coli* (C3040, NEB), except for *pBAD* promoter mutagenesis where transformations were performed directly on attenuated *Salmonella enterica serovar typhimurium SL1344* (Stm; a gift from Jeff Hasty, University of California, San Diego). All constructs were verified via sequencing.

### Manufacturing Dual-Layer 96-Well Agar Plates

Lennox Luria Broth (LB) agar was prepared, autoclaved and allowed to cool to 65°C before any reagents were added to it. Wells were first loaded with 125 µl of agar containing the relevant induction reagents (IPTG, L-arabinose) and antibiotics, and incubated at room temperature for ∼10 minutes for this first layer to polymerize. The suppression layer was then loaded into each well as a second 125-µl volume, this time containing the relevant suppression reagents (D-glucose) and antibiotics. The second layer was allowed to polymerize and cool to ≤37°C before bacteria patching. Plates were prepared within 2 hours of bacteria patching.

### Cell Culture and Cell Line Generation

BL21-AI *E. coli* was acquired from Invitrogen (C607003), attenuated *Salmonella enterica serovar typhimurium SL1344* (Stm) was a gift from Jeff Hasty (University of California, San Diego), and *E. coli Nissle 1917* (EcN) was acquired from Mutaflor®. Unless otherwise stated, all plasmids were introduced into bacteria via electroporation, followed immediately by addition of SOC media containing 0.5% D-glucose (m/v) and incubation for 2-3 h at 250 rpm, 37°C before introduction of any antibiotics. Cells were then inoculated in Lennox LB containing 0.5% D-glucose (m/v) with antibiotics and grown at 250 rpm, 37°C for 6-8 h to reach log phase ahead of patching on 96-well dual-layer agar plates, or on 10-cm dual-layer agar plates.

### Cell Patching for Solid Phase Expression

For patching, 5-8 µl of bacteria were dispensed onto agar, and allowed to incubate at 37°C for 24 h to allow for gene expression. Bacteria exhibited significantly higher patch opacity in cases of successful GV assembly. Cells were then collected by dispensing 100 µl of PBS and incubating at room temperature for ∼10 minutes, before mixing the sample and collecting it into a secondary apparatus for downstream measurements. Bacteria exhibiting successful GV assembly should start to float in PBS within the 10-minute incubation period. GV assembly was induced on solid-phase agar in 96-well format unless otherwise stated.

### GvpC Library Construction and Expression

All PCR reactions were performed using the Taq Hot Start System (M0496, NEB) and fragment insertion was conducted with the High Fidelity DNA Assembly System (M0541, NEB). Error prone PCR of the *gvpC* sequence was performed by adding 20 to 250 μM MnCl_2_ to the reaction and amplifying across 30 cycles. PCR products were then cloned into the ΔgvpC plasmid backbone, replacing *gfp*, and electroporated into BL21-AI *E. coli* (C607003, Invitrogen). The pooled bacteria population was spread onto 10-cm dual-layer agar plates containing 60 µM IPTG, 0.3% L-arabinose (m/v), 0.5% D-glucose (m/v) and 2× kanamycin, and incubated overnight at 37°C. Single white colonies were picked to inoculate 1.5-ml volumes of Lennox LB containing 2% D-glucose and 2× kanamycin and grown at 250 rpm, 37°C for 6-8 hours, prior to being patched onto 96-well dual-layer agar plates containing 60 µM IPTG, 0.3% L-arabinose (m/v), 0.5% D-glucose (m/v) and 2× kanamycin, and incubated at 37°C for 24 hours for expression.

### Ultrasound Imaging

All agarose solutions used for ultrasound imaging were passively degassed for at least 24 h before casting. Cells suspended in PBS were mixed 1:1 (v/v) with 1% low-melt agarose (m/v) and loaded into imaging phantoms made of 2% agarose (m/v). Imaging was performed using a Verasonics Vantage programmable ultrasound scanning system and a L22-14vX 128-element linear array Verasonics transducer, with a specified pitch of 0.1 mm, an elevation focus of 8 mm, an elevation aperture of 1.5 mm and a center frequency of 18.5 MHz with a 67% -6 dB bandwidth. All image acquisition scripts were coded on MATLAB (Mathworks). *In vitro* samples were centered at a depth of 5 mm during acquisition. For nonlinear imaging, a custom X-wave amplitude modulation (xAM) sequence with a cross angle (θ) of 19.5°, an aperture of 65 elements, and a transmit frequency of 15.625 MHz was used. Each image was an average of 30 accumulations. For linear image acquisition, a conventional ray-line scanning B-mode pulse sequence with parabolic focusing at 10 mm and an aperture of 40 elements was used. A parabolic B-mode pulse set at the maximum transmit voltage (30V) was used to collapse GVs. The transmitted pressure at the sample position at 5 mm was measured using a Precision Acoustics fiber-optic hydrophone system.

### Linear Unmixing of Acoustic Signals

All image processing scripts were coded on MATLAB (Mathworks). To generate the multiplexing matrix ‘*M*’, reference samples of Stm encoding bARG_560_ and EcN encoding bARG_710_ were imaged using the xAM pulse sequence across the pressure domain. Regions-of-interest were segmented for each sample and organized into an *N*×*P* matrix, where *N* is the number of acoustic colors, *P* is the number of pressure acquisitions, and the value in each cell represents the xAM signal for a given color at a given pressure. Once *M* was established, all unknown images thereafter were acquired following the same acquisition parameters as the reference samples. Unknown images were first thresholded (∼100 a.u. for *in vitro* samples, ∼200 a.u. for *in vivo* images), such that if the nonlinear signal of a pixel did not surpass this threshold at any pressure acquired, it was zeroed out on the image, and by extension, on the coefficient matrix ‘*C*’. Linear unmixing was then performed pixelwise to solve for the coefficient matrix ‘*C*’ to generate a matrix of size *Z*×*X*×*N*, where *Z* is the depth and *X* is the width of the deconvoluted image. The following equation was used to solve for *C*, using a non-negative least squares (NNLS) fitting, where λ is the nonlinear signal as a function of pressure for each pixel:

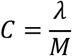

### Mammalian Cell Culture

Murine colon carcinoma cells (MC26) were acquired (Cat. 400156, BioHippo) and grown in Dulbecco’s modified Eagle’s medium supplemented with 10% fetal bovine serum (FBS) and 1× antibiotic-antimycotic.

### Animal Models

All animal protocols were approved by the Institutional Animal Care and Use Committee (IACUC) at the California Institute of Technology (Protocol #1735) and comply with federal and state regulations governing the humane care and use of laboratory animals. Animals were housed in a facility maintained at 21–24°C and 30–70% humidity, with a lighting cycle of 13 hours on (6:00– 19:00) and 11 hours off. Female BALB/c mice (Strain #000651, Jackson Laboratory) were used for all animal experiments. Mice were anesthetized with 1–3% isoflurane in 100% oxygen using a nose cone co-linked to a vacuum line for active scavenging, and kept warm on a heated stage.

### Ultrasound Imaging of the Gastrointestinal Tract

For imaging the gastrointestinal tract, mice were placed in the supine position, depilated over the imaging region, and received an enema with PBS to clear the tract of feces using a flexible plastic tubing needle tip (Cat. CAD9918, Millipore Sigma). Prior to imaging, Stm encoding bARG_560_ and EcN encoding bARG_710_ were induced for GV expression as described above, collected and diluted in pure or mixed populations (1:1 v/v) to a concentration of 2×10^8^ cells ml^-1^, and then mixed in a 1:1 ratio with 42°C 1% low-melt agarose. An 8-gauge needle was filled with the gelled bacteria solution and incubated at room temperature to polymerize. The bacteria-embedded gel was then injected into the colon of the mouse with a PBS back-filled syringe. B-mode imaging was used to identify the colon lumen. Nonlinear ultrasound imaging was then performed as described above.

To ensure that the nonlinear signals we were observing arose from GVs and not from the interface of the gastrointestinal tract with its lumen and/or extraluminal fluid of the peritoneal cavity, we also applied a parabolic B-mode collapse pulse to irreversibly collapse the GVs and acquired post-collapse xAM images (**Supplementary Fig. S7**). Following acquisition, we processed the raw image series by filtering out pixels that did not generate significant nonlinear signals at any point throughout the acquisition series (S < 200 a.u.). In some cases, we also discarded single pressure acquisitions from the image series because of non-negligible movement from the mouse during breathing. For analysis, we segmented regions-of-interest representing the gastrointestinal lumen from the raw nonlinear image, which was then overlaid on the coefficient matrices to quantify mean signal values from each acoustic channel.

### *In Vivo* ARG Expression by Tumor-Colonizing Bacteria

Mice were injected subcutaneously at the right flank with 3×10^5^ MC26 cells and solid tumors formed over 2-3 weeks (200–400 mm^3^). The day before injection of bacteria, ibuprofen was added to the drinking water at 0.2 mg ml^−1^. Stm encoding bARG_560_ and EcN encoding bARG_710_ were inoculated in Lennox LB containing 0.5% D-glucose (m/v) with antibiotics and grown at 250 rpm, 30°C for 12-16 h. Cells were then washed 3× with PBS and diluted to OD_500_ = 0.300. 150 µl of single- or dual-species bacteria solutions were then injected intravenously into the tail veins of mice. Cells were given 24 h following intravenous injection to colonize tumors, after which mice were injected intraperitoneally every 24 h for 3 days with 120 mg of L-arabinose to induce expression of bARG_560_ and/or bARG_710_. On the fourth day, mice were placed in a prone position, depilated over the imaging region, and nonlinear ultrasound imaging was performed on a cross-section of the tumor. Mice were sacrificed immediately after ultrasound imaging for histology.

### Histology

Whole-tumors were excised from mice immediately after ultrasound imaging and placed in 4% paraformaldehyde for at least 48 h. Following fixation, tumors were moved into 30% sucrose (m/v) for at least 24 h, or until they sank to the bottom of the solution. Tumors were dried for approximately 30 minutes with paper towel, positioned in a volume of OCT freezing medium and incubated at -30°C overnight. Tumors were then cut into 20-µm thick sections and placed onto microscope slides using a Leica CM1950 cryostat. Fluoro-gel Mounting Medium (Cat. 17985-50, Electron Microscopy Sciences) was applied dropwise to tumor sections, and then encased with glass coverslips. Slides were imaged with a Zeiss LSM 800 Confocal Laser Scanning Microscope with a 5x objective for whole-tumor tiled imaging, and 20x objective for localized imaging of the tumor core and tumor rim.

### Statistical Analysis

No statistical methods were used to predetermine sample size. Error bars indicate the standard error of the mean (s.e.m.). A threshold of 0.05 was used for statistical significance. P values were calculated using a two-tailed paired t-test, an ordinary 1-way analysis of variance (ANOVA), or a 2-way ANOVA.

## Data and Materials Availability

Plasmids used in this study will be available from M.G.S. under a material agreement with the California Institute of Technology at the time of manuscript publication. Processing scripts used to generate key figures and results will be posted to a publicly accessible repository at the time of manuscript publication. Raw data can be made available by M.G.S. upon reasonable request.

## Acknowledgements

The authors thank M. Buss for her advisement in microbial protocols and E. Criado-Hidalgo for hydrophone measurements. This research was supported by the Chan-Zuckerberg Initiative, and the National Institutes of Health (R01-EB018975 to M.G.S.). N.N.N. is grateful to have received support as an Amgen awardee of the Life Sciences Research Foundation, and support from the Caltech Center for Environmental Microbial Interactions. M.G.S. is an investigator of the Howard Hughes Medical Institute.

## SUPPLEMENTARY INFORMATION

**Supplementary Note 1 - Structural and mechanistic predictions of GvpC**_**L154P**_ Because of the restricted rotation of its N-C_α_ bond, and lack of hydrogen atom for hydrogen bonding, we expected the proline to disrupt the alpha helical structure of GvpC in some way. We processed the mutated sequence using AlphaFold to generate a structural prediction with a confidence score (pLDDT) of 76.9%. First, we found that L154 was part of the shell-binding face of GvpC, resulting in direct loss of a single shell-binding residue. Additionally, L154P generated a significant structural kink that rigidly deviates the final 21% length of the GvpC helix away from its binding interface on the shell (Δα = 12.1° relative to wild-type; **Supplementary Fig. S3**)

For example, the next shell-binding residue after L154P, F161, was situated 2.1 Å from its original alignment, which is enough to weaken, if not disrupt, its speculated interaction with E60 of gvpA1 (**Supplementary Fig. S3**).

Binding residues further downstream were increasingly misaligned. These measurements were made on the free-form structure of GvpC, which may differ from analysis performed on a shell-bound structural analysis of GvpC, but because knowledge of the binding mechanisms between the GvpA shell protein and the GvpC scaffolding protein is still limited, it is unlikely that this type of analysis will yield any more tangible insight at this point.

**Supplementary Note 2 - Achieving bARG**_**560**_ **and bARG**_**710**_ **expression in probiotic strains** We transitioned our constructs from T7-dependent circuits to the directly inducible *pBAD* promoter, but initially observed no GV assembly from either construct (bARG_560_, bARG_710_) in either strain (Stm, EcN) across a range of induction conditions. First, we reasoned that removal of genes not directly contributing to GV assembly could reduce burden and increase GV yield through reallocation of finite transcriptional and translational resources.

We tested single gene deletions of *gvpW, gvpX, gvpY, gvpH*, and *gvpZ* in BL21-AI *E. coli*, as these were already deemed nonessential for GV assembly in the native organism, in addition to deletions of *gvrB* and *gvrC*, as these were putative regulatory proteins that would be obsoleted by our synthetic control circuit. We saw GV yield increase across all of our single gene deletion constructs relative to the original wild-type construct (bARG_ser_), in agreement with our finite resources hypothesis (**Supplementary Fig. S4**).

We then tested a stepwise gene deletion tree in EcN, starting with bARG_560_ as the parent construct (*gfp*^+^, Δg*vpC*), and found that combined deletions of *gvpW, gvrB* and *gvrC* enabled GV assembly in this microbe across a range of induction conditions (**Supplementary Fig. S5**). Next, we moved this new, smaller bARG_560_ construct into Stm, but again observed no assembly. We then targeted the I1 and I2 operators of *pBAD* for mutagenesis, and found a mutant that enabled GV assembly from bARG_560_ in Stm (**Supplementary Fig. S5**).

## SUPPLEMENTARY FIGURES

**Supplementary Figure S1.**
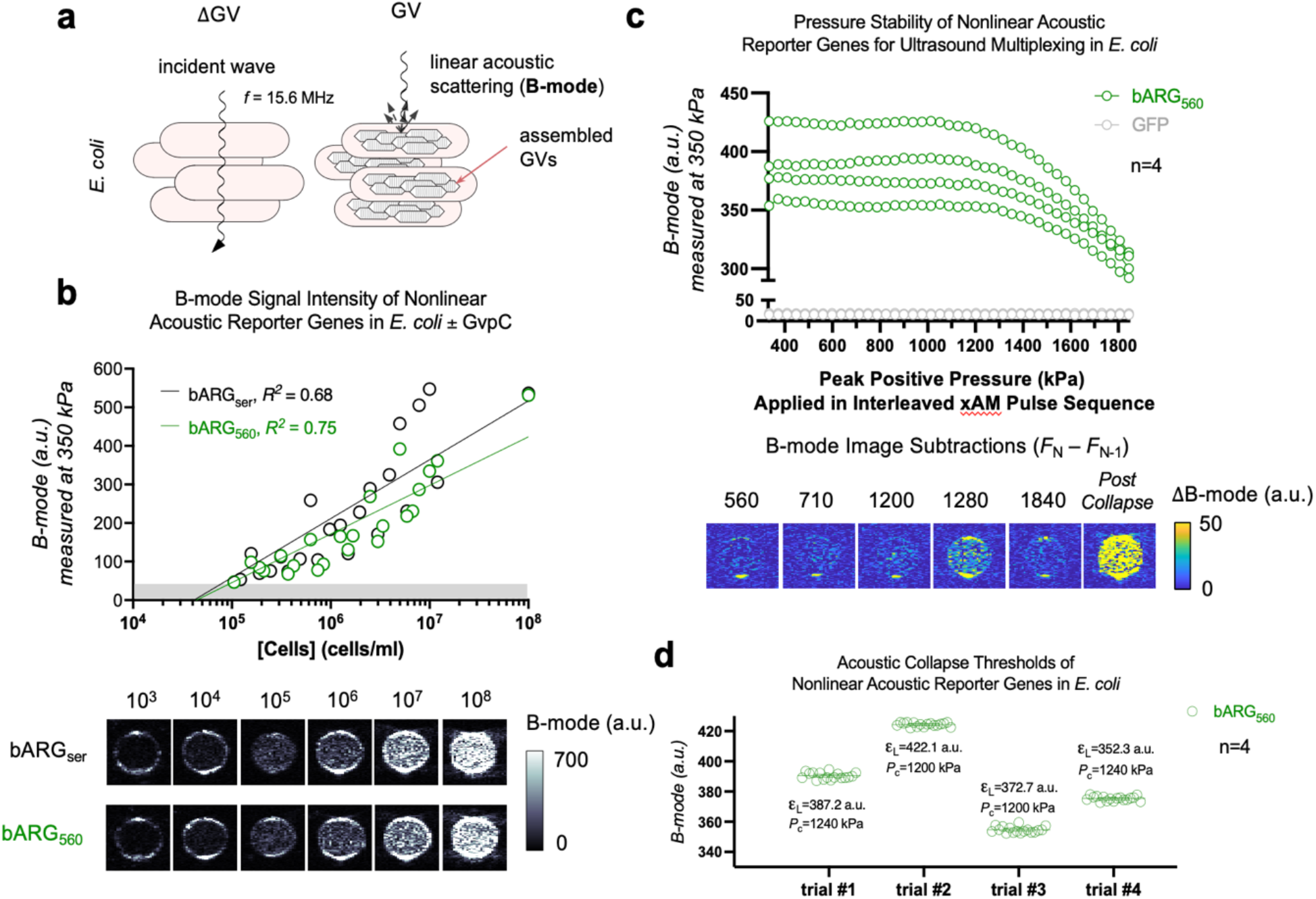
Determination of the acoustic collapse pressure of bARG_560_. **(a)** *E. coli* inherently produce minimal linear scattering on ultrasound, but GVs produced within *E. coli* can operate as linear scattering agents at low acoustic pressures on B-mode images. **(b)** B-mode signal intensity (a.u.) at 350 kPa measured across increasing concentrations of *E. coli* (10^3^– 10^8^ cells ml^-1^, n=3) encoding bARG_Ser_ (*black*) or bARG_560_ (*green*), and representative B-mode images of each. **(c)** B-mode signal intensity (a.u.) at 350 kPa of *E. coli* encoding bARG_560_ (n=4) following X-wave amplitude modulation (xAM) pulse sequence imaging at increasing pressures (0.3–1.8 MPa, Δ*P*_N-(N-1)_ ≅ 40 kPa). B-mode image subtractions (a.u.). The “Post Collapse” image represents the subtraction between a B-mode image acquired after applying the maximum xAM pulse sequence pressure (1.8 MPa) and a B-mode image acquired after applying a parabolic B-mode collapse pulse. **(d)** To determine the acoustic collapse pressure of bARG_560_ nanostructures, B-mode signal intensity at 350 kPa was averaged across 10 acquisitions for each sample, and the lower limit of the 95% confidence interval (ε_L_) was calculated for each trial (n=4). The collapse pressure was defined as the threshold at which B-mode signal intensity first fell below the lower limit of the 95% confidence interval.

**Supplementary Figure S2.**
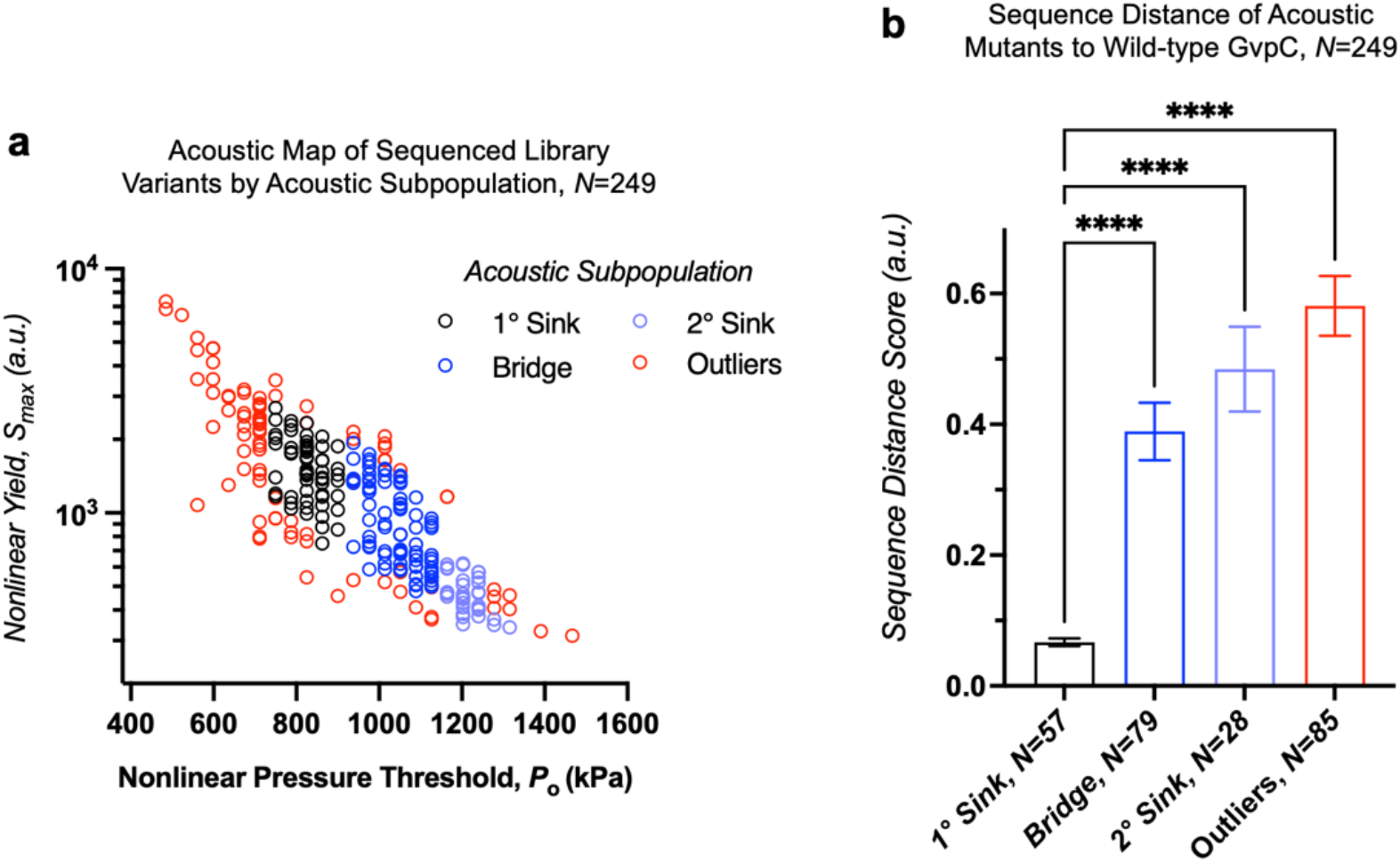
Sequence distance scores of mutants to wild-type *gvpC* by acoustic subpopulation. **(a)** Acoustic map of sequenced library variants according to their acoustic subpopulation: 1° acoustic sink centered on the wild-type *gvpC* sequence (*black)*, 2° acoustic sink (*purple*), the bridge between them (*blue*), and any mutants that fell outside of these regions (*red*). Refer to Fig. 2b for the parent acoustic map containing all variants (sequenced and not sequenced), colored by the relative density of mutants. **(b)** Average sequence distance scores (mean ± s.e.m., a.u.) of acoustic subpopulations following Needleman-Wunsch alignment to wild-type *gvpC*. ****p<0.0001.

**Supplementary Figure S3.**
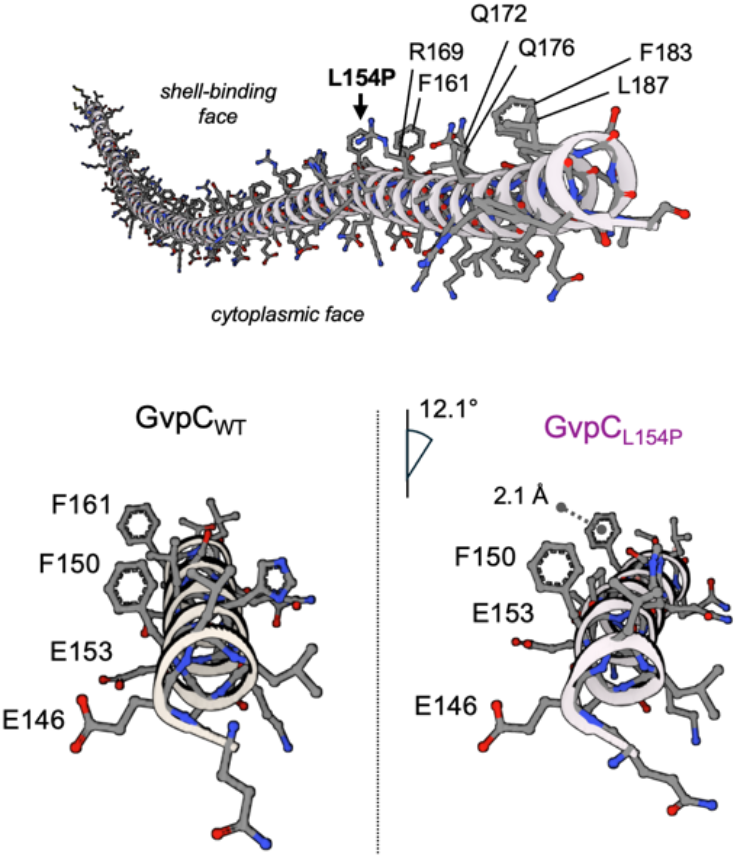
AlphaFold structural prediction of GvpC_L154P_. *Top:* Ball-and-stick model of the AlphaFold structural prediction of GvpC_L154P_ with GV-binding residues labelled downstream of the L154P structural kink. *Bottom:* wild-type gvpC structure (*left*) and GvpC_L154P_ (*right*) spanning from Q145 to T165, with angle deviation (12.1°) and F161 distance differential (2.1 Å) denoted.

**Supplementary Figure S4.**
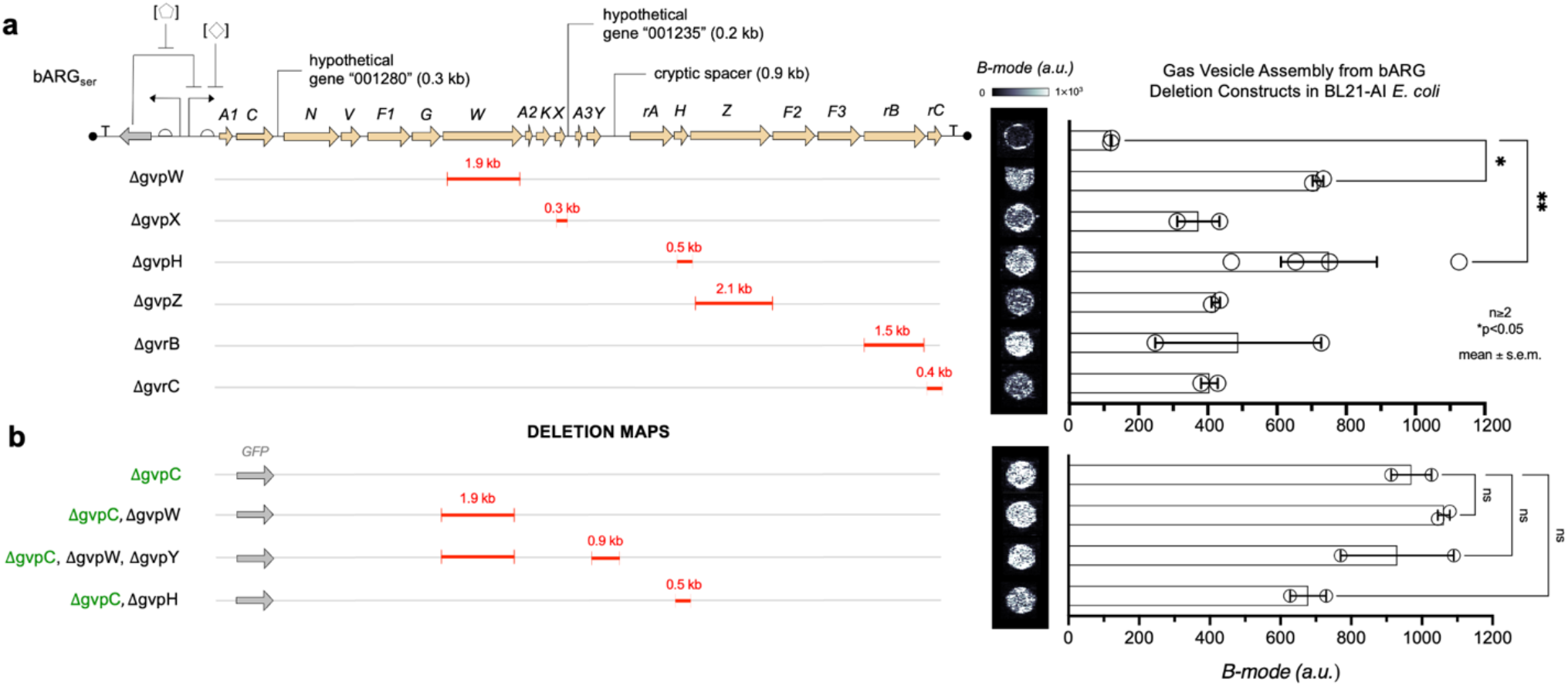
GV assembly screening of gene deletion constructs in BL21-AI *E. coli*. **(a)** Construct and single gene deletion maps based on the bARG_Ser_ construct. B-mode signal intensity (a.u.) at 350 kPa of bARG_Ser_ and single gene deletion constructs measured from 10^6^ cells ml^-1^ in BL21-AI *E. coli* (mean ± s.e.m., n≥2). **(b)** Multi-gene deletion maps based on the bARG_560_ construct. B-mode signal intensity (a.u.) at 350 kPa of bARG_560_ and multi-gene deletion constructs measured from 10^9^ cells ml^-1^ in BL21-AI *E. coli* (mean ± s.e.m., n≥2).

**Supplementary Figure S5.**
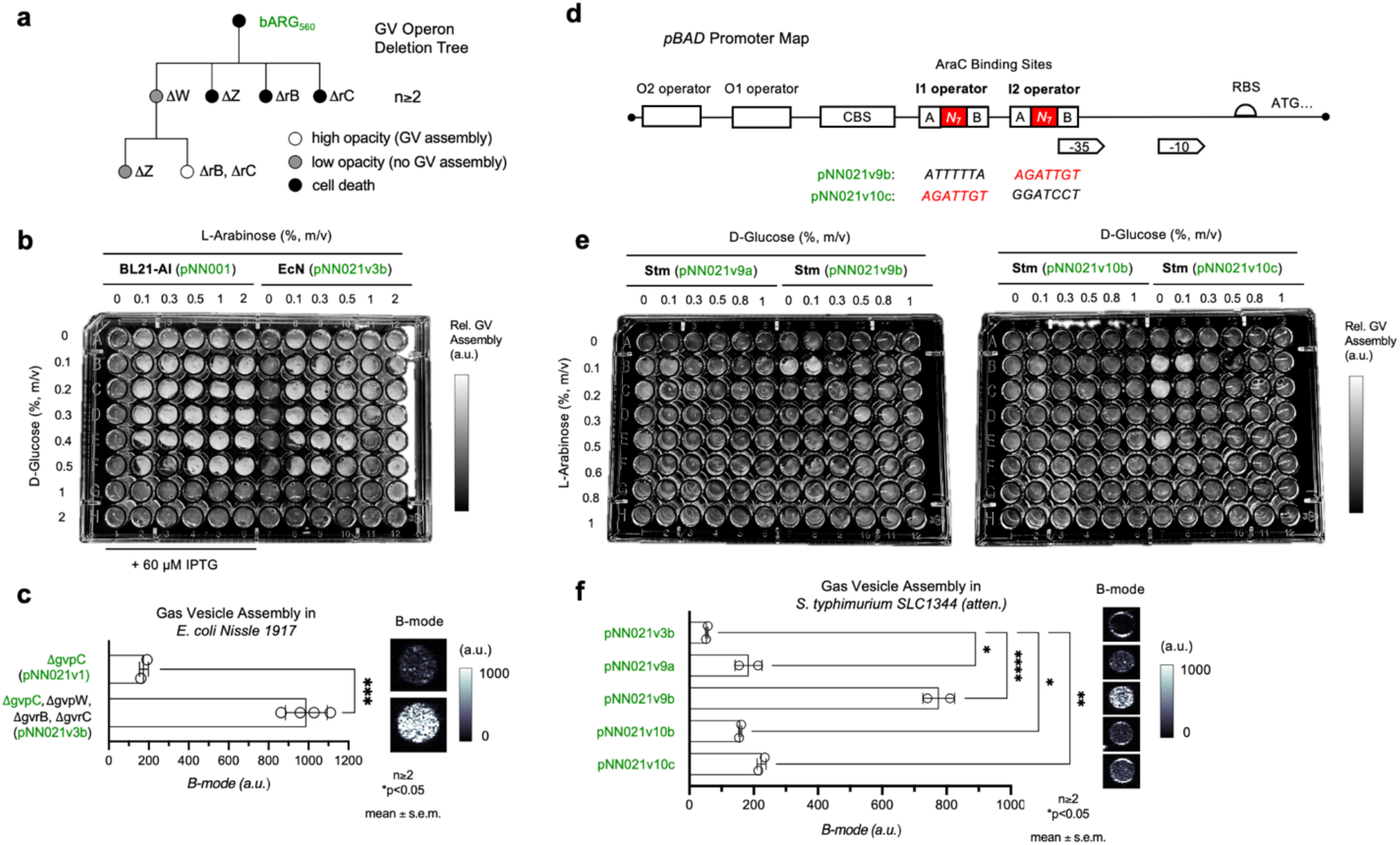
Engineering GV assembly constructs for probiotic microorganisms. **(a)** Synthetic evolutionary tree of bARG_560_ in *E. coli Nissle 1917* (EcN). Each construct was transformed into EcN and tested across a range of induction conditions, which resulted in either cell death upon induction (*black*), or cell survival with (*white*) or without (*grey*) GV assembly (n≥2). **(b)** Representative 96-well patch plate of BL21-AI *E. coli* transformed with bARG_560_ in a T7-circuit (pNN001; *left half of plate*) and *E. coli Nissle 1917* (EcN) transformed with bARG_560_ that includes deletions of gvpW, gvrB, and gvrC (pNN021v3b; *right half of plate*), induced for GV expression on dual-layer agar containing variable concentrations of L-arabinose (0–2%, m/v) and D-glucose (0–2%, m/v). Non-variable conditions include 60 µM IPTG for BL21-AI *E. coli*, 37°C incubation, and 5 g/L NaCl. **(c)** B-mode signal intensity (a.u.) at 350 kPa of *E. coli Nissle 1917* transformed with pNN021v1 (pBAD, *gfp*^+^, Δg*vpC*) or pNN021v3b (pBAD, *gfp*^+^, Δg*vpC*, Δg*vpW*, Δg*vrB*, Δg*vrC;* mean ± s.e.m., n≥2). **(d)** Promoter map of pBAD. AraC binding sites include the O2, I1, and I2 operators. CBS, cAMP receptor protein (CRP) binding site. RNA polymerase binding sites, -35 and -10 elements. RBS, ribosome binding site. ATG, start codon. The 7 nucleotides within the I1 and I2 operators, internal to the A and B boxes, were targeted for random mutagenesis in attenuated *Salmonella typhimurium SL1344* (Stm). Successful mutant sequences from the pooled library that exhibited GV assembly (pNN021v09b, pNN021v10c) are included: wild-type sequence in black, mutant sequence in red. **(e)** Representative 96-well patch plate of select Stm colonies from a pooled library transformation of I1 and/or I2 mutant constructs, induced for GV expression on dual-layer agar containing variable concentrations of L-arabinose (0–1%, m/v) and D-glucose (0–1%, m/v). pNN021v09b and pNN021v10c exhibited increased opacity, indicative of GV assembly. **(f)** B-mode signal intensity (a.u.) at 350 kPa of Stm transformed with pNN021v3b (pBAD, *gfp*^+^, Δg*vpC*, Δg*vpW*, Δg*vrB*, Δg*vrC*) and various pBAD mutant constructs derived from pNN021v3b (mean ± s.e.m., n≥2).

**Supplementary Figure S6.**
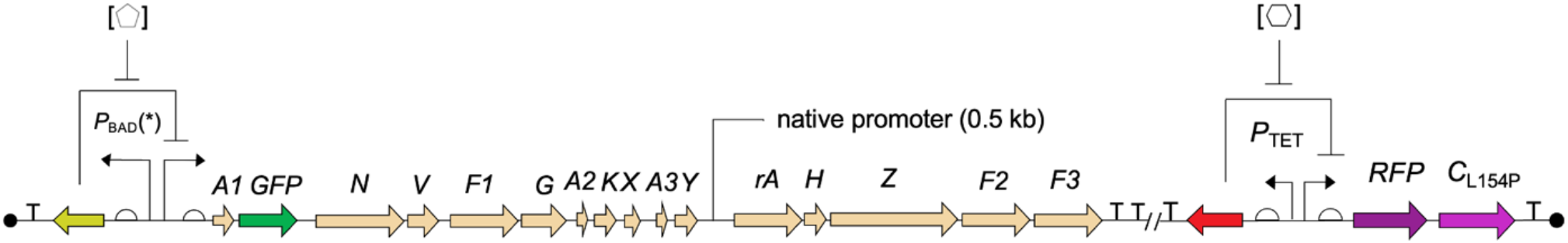
The minimized bARG_560_ operon and the tetracycline-responsive gvpC_L154P_ operon for cell state imaging. The chemical inducer used for the *pBAD*(*) promoter is L-arabinose and the chemical inducer for the *pTET* promoter is anhydrous tetracycline (aTc).

**Supplementary Figure S7.**
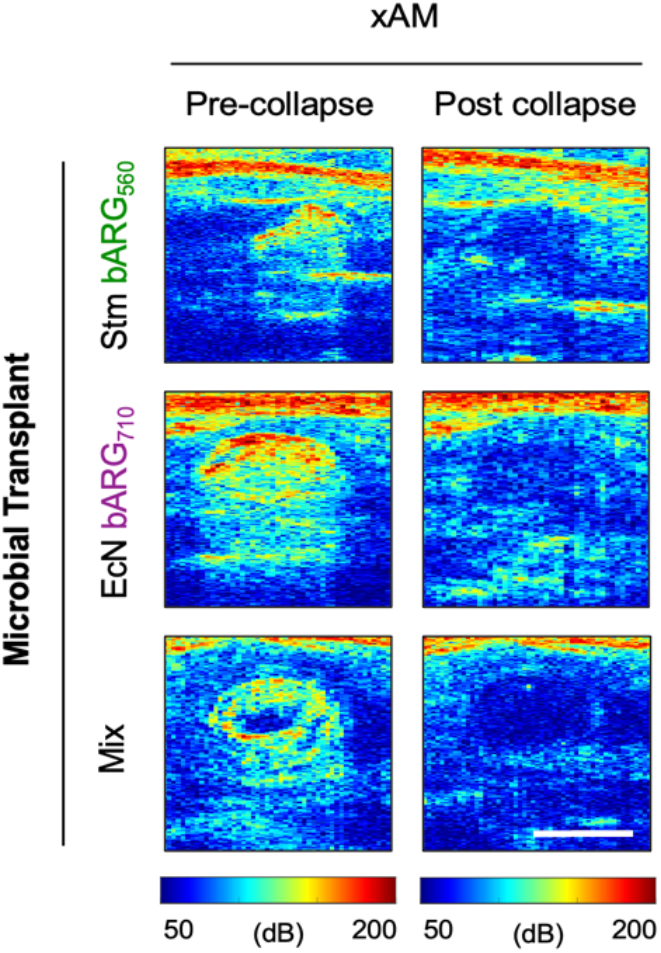
Nonlinear signal validation in the gastrointestinal tract of live mice. Representative X-wave amplitude modulation (xAM) images at ∼1000 kPa of the gastrointestinal system of mice transplanted with attenuated *S. typhimurium SL1344* (Stm) encoding bARG_560_ and/or *E. coli Nissle 1917* (EcN) encoding bARG_710_, before and after application of a high-pressure collapse pulse. Decibels, dB. Scale bar, 2 mm.

**Supplementary Table T1.** Circuit variables and parameter limits for GV expression by bARG_560_.

| Variable |  | Parameter Limits for GV Assembly |
| --- | --- | --- |
| 1 | bARG Operon T7 RNA Polymerase-dependent Promoter Circuit | pLac |
| 2 | bARG Operon Inducer Concentration | IPTG, 40–70 μM |
| 3 | T7 RNA Polymerase Operon Suppressor Concentration | D-glucose, $\leq 0.5\%$ (m/v) |
| 4 | T7 RNA Polymerase Operon Promoter Circuit (Expression Strain Background) | pBAD (BL21-AI<sup>TM</sup>, cat no. C607003) |
| 5 | T7 RNA Polymerase Operon Inducer Concentration | L-arabinose, 0.3–0.5% (m/v) |
| 6 | NaCl Concentration | 5–10 g/L |
| 7 | Induction Temperature | 30–37°C |

